# Directed evolution of a sequence-specific covalent protein tag for RNA labeling

**DOI:** 10.1101/2024.08.30.610577

**Authors:** Rongbing Huang, Alice Y. Ting

## Abstract

Efficient methods for conjugating proteins to RNA are needed for RNA delivery, imaging, editing, interactome mapping, as well as for barcoding applications. Non-covalent coupling strategies using viral RNA binding proteins such as MCP have been applied extensively but are limited by tag size, sensitivity, and dissociation over time. We took inspiration from a sequence-specific, covalent protein-DNA conjugation method based on the Rep nickase of a porcine circovirus called “HUH tag”. Though wild-type HUH protein has no detectable activity towards an RNA probe, we engineered an RNA-reactive variant, called rHUH, through 7 generations of yeast display-based directed evolution. Our 13.4 kD rHUH has 12 mutations relative to HUH, and forms a covalent tyrosine-phosphate ester linkage with a 10-nucleotide RNA recognition sequence (“rRS”) within minutes. We engineered the sensitivity down to 1 nM of target RNA, shifted the metal ion requirement from Mn^2+^ towards Mg^2+^, and demonstrated efficient labeling in mammalian cell lysate. This work paves the way toward a new methodology for sequence-specific covalent protein-RNA conjugation in biological systems.

## Introduction

RNAs play a central role in encoding, regulating, and driving biological function. Increasingly, RNA-based pathways are also being intercepted by therapeutic modalities to treat disease(1). Due to their central importance, technologies to detect, image, manipulate, and engineer RNAs are in great demand. However, many of the most robust molecular tools, especially for deployment in complex biological settings such as living cells, are encoded as proteins, not RNA. These include fluorescent proteins, CRISPR/Cas enzymes, HaloTag(2), light-switchable proteins such as LOV and CRY/CIBN, endonucleases, viral proteases, and the proximity labeling enzymes APEX(3) and TurboID(4). To use these tools, it is often necessary to establish a molecular link between protein and RNA, to bring protein-encoded function to specific RNAs of interest.

It is possible to link proteins and RNA together using conjugation chemistry, but cost, yield, imperfect specificity, and the need for subsequent purification can be prohibitive. Genetically-encoded strategies for joining RNA to proteins provide attractive alternatives due to their ease of use and compatibility with living systems. Most common among these are the MS2/MCP system(5, 6), boxB/lambdaN(7), and PP7/PCP(8). For example, the MS2/MCP system consists of an MS2 hairpin (19 nt) fused to an RNA of interest, which recruits MCP protein (13.7 kD) fused to a protein of interest. This versatile methodology has been used to image cellular RNAs(6, 8, 9), deliver transcriptional regulators to sgRNAs(10), target proximity labeling enzymes to RNA for discovery of interaction partners(11) and facilitate RNA editing(12). However, a major limitation of these sequence-specific RNA binding proteins is that the linkages are non-covalent. The pairs dissociate over time and cannot withstand extreme conditions such as higher temperature, salt, pH, or organic solvent. A genetically-encoded but covalent methodology for linking specific proteins and RNA together would offer a new paradigm.

To develop such a method, we looked for modular systems that provide covalent RNA-protein conjugation, are fully genetically-encoded (no need for chemical derivatization or post-transcriptional/translational modifications), fast, and sequence-specific. We did not identify a method that met all these criteria, but found that the HUH tag system(13) provides an analogous methodology for covalently coupling single stranded DNA (ssDNA) to protein. Derived from a porcine circovirus (PCV2) Rep nickase domain, the HUH tag (herein referred to as dHUH) recognizes a 10 nt ssDNA recognition sequence (herein referred to as dRS for DNA recognition sequence) and forms a covalent adduct via nucleophilic attack of Tyr96 of dHUH onto a specific phosphate backbone linkage of dRS. The outstanding specificity, speed, and sensitivity of this coupling has enabled the HUH tag system to be used for a wide variety of applications, including conjugation of ssDNA barcodes to antibody reagents, tethering template DNA to Cas9 for genome editing(14), and displaying DNA on nanoparticles(15).

Due to the similarities between ssDNA and RNA, we wondered whether it would be possible to engineer dHUH to recognize and form covalent adducts with short sequence-specific RNA tags (“rRS”) instead of DNA (**Figure 1A**). Herein we describe the engineering of “rHUH” protein for covalent coupling to short (10 nt) RNA motifs (rRS) and the characterization of this new conjugation system on yeast and in vitro.

**Figure 1.**
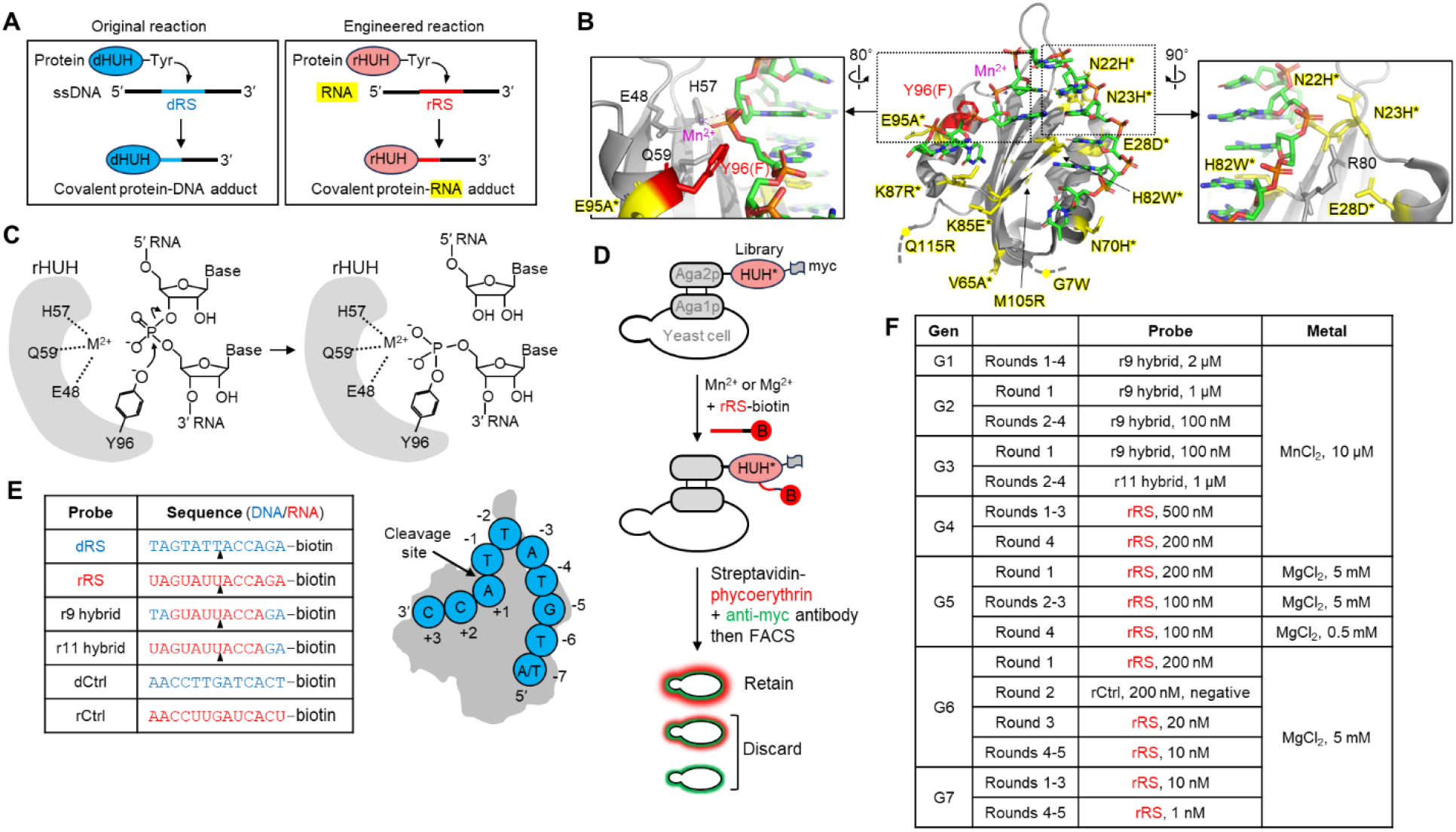
Engineering rHUH and directed evolution scheme. (**A**) Schematic of wild-type reaction of dHUH protein with ssDNA (the DNA recognition sequence, dRS), and desired conjugation between engineered rHUH protein and ssRNA (the RNA recognition sequence, rRS). (**B**) Crystal structure of wild-type HUH protein in complex with its target ssDNA sequence (dRS), from PBD ID 6WDZ [ref]. The nucleophilic Tyr96 (mutated to Phe in this structure) is red, the Mn^2+^ atom is purple, and the 12 residues mutated over the course of evolution to rHUH are colored yellow. Zoom on right shows multiple mutations proximal to -3 and -4 positions of bound dRS. *Indicates mutation found in clone G5. G7 contained all the shown mutations. (**C**) Expected active site chemistry of rHUH, showing nucleophilic attack by deprotonated Tyr96 on RNA backbone, leading to a covalent adduct between rHUH protein and the 3’ portion of the target RNA (rRS). M^2+^, divalent metal ion such as Mn^2+^ or Mg^2+^. (**D**) Yeast surface display selection scheme. To a library of HUH mutants (HUH*) is added biotinylated rRS or hybrids. After washing, cells are stained with streptavidin-phycoerythrin (PE) to quantify probe binding and anti-myc antibody to quantify HUH expression level. FACS sorting is used to enrich cells with high streptavidin-PE/myc intensity ratio. (**E**) Sequence of ssDNA target sequence (dRS), ssRNA target sequence (rRS), and RNA-DNA hybrids used for early generations of directed evolution. (**F**) Table of selection conditions used across 7 generations of directed evolution. Each generation consisted of 4-5 rounds of amplification and sorting, followed by identification of the top clone (named “G1-G7”) and re-diversification via error-prone PCR.

## Results

The structure of dHUH in complex with dRS(16) shows extensive contacts between the protein and both phosphate backbone and nucleobase features of dRS, which is bent into a U-shape with hydrogen bonding between two base pairs at the hairpin turn (**Figure 1B**). The active site of dHUH contains a Mn^2+^ ion, coordinated by Glu48, His57 and Gln59 sidechains, which helps to stabilize the pentavalent phosphate transition state generated by nucleophilic attack from Tyr96 (**Figure 1C**). After the reaction, the 3’ end of dRS is covalently coupled to dHUH via a tyrosine-phosphate ester bond, while the 5’ end of dRS (7 nt long) is released.

We established a yeast surface display platform to both characterize and evolve dHUH (**Figure 1D**). The 13.3 kD protein was fused to the C-terminal end of the yeast mating protein Aga2p for display on the cell surface. A myc tag was fused to the C-terminal end of dHUH to facilitate detection by anti-myc antibody. To assess the activity of displayed dHUH, we added 3’-biotin-conjugated DNA probe (biotin-dRS, **Figure 1E**) to the cells, washed, and stained with streptavidin-phycoerythrin conjugate to assess labeling. We found robust labeling of dHUH by biotin-dRS down to a concentration of 0.1 nM, but no labeling by a scrambled control DNA probe (**Figure S1A**).

We then tested the RNA version of the dRS probe (biotin-rRS). No signal was detected, even at high (1 uM) probe concentrations and long incubation times (2 hours). We attempted some rational mutagenesis of dHUH, based on the structure, selecting sidechains that might contact the 2’-OH of bound RNA (**Figure S1B**). None of the 7 mutants we tested showed activity with RNA probe, though some had decreased reactivity towards DNA. We recognized that much more extensive screening would be necessary and proceeded to build a library of dHUH variants for yeast display. To construct the library, we opted for error prone PCR, rather than more focused mutagenesis of residues in contact with dRS, because long-range interactions could impact both substrate recognition and catalysis. Furthermore, we did not assume that a rRS probe would dock to HUH protein in the same manner as dRS. Our error prone PCR produced an average of 1-2 amino acid changes per gene.

We were concerned that the change from dRS to rRS would be too drastic, such that no member of our initial library might have the ability to recognize a fully RNA substrate. Thus, we performed a study of DNA-RNA hybrid substrates, in which single nucleotides or groups of nucleotides in dRS were replaced with RNA (**Figures S1C-D**). Our results indicate that several positions of dRS are tolerant to substitution by RNA, while the -3 and -4 positions are quite sensitive (**Figure S1E**). Ultimately, we selected an r9 hybrid probe (**Figure 1E**), because we could detect labeling of wild-type dHUH at high concentrations of this probe (**Figure S1F**).

We began evolution by treating the yeast-displayed dHUH library with 2 uM of r9 hybrid for 1 hour. After washing the cells and staining with streptavidin-PE and anti-myc antibody, we found that ∼30% of myc-positive cells showed signal above background. We used FACS to separate this population, amplified the yeast cells, and performed 3 more rounds of selection. Enriched cells were sequenced, and unique clones were compared on the yeast cell surface (**Figure S2**). The best clone, with 3 mutations relative to dHUH, was named G1 and used as the template for the second generation of evolution.

In this manner, we performed 7 total generations of directed evolution, with the winning clone from the previous generation serving as the template for library generation for the next generation (**Figure 1F**). Generation 2 still utilized the r9 hybrid probe, while Generation 3 used an r11 hybrid with only two nucleotides from DNA. By Generation 4, we had sufficient activity to switch to an entirely RNA-based probe. In Generations 4-7, we progressively decreased the concentration of rRS (from 500 nM to 10 nM) to select for HUH variants with higher RNA affinity. In Generation 5, we also switched from MnCl_2_ supplementation to MgCl_2_ supplementation, as the latter but not former is available in cells.

## Results from directed evolution

The sequences of the winning clones from each generation (G1-G7) are shown in **Figure S1A**. On yeast, we compared the clones G1-G5 alongside the original dHUH for reaction with both dRS and rRS. **Figure 2A** shows that labeling with rRS is detectable at G3 and jumps dramatically at G4. Reaction with dRS drops somewhat over generations with the most dramatic dip at G3, perhaps due to the acquisition of the H82Y mutation, which may confer strong RNA-bias. Unfortunately, G4 and G5 also showed significant reactivity towards a scrambled rRS control sequence (“rCtrl”), although the labeling extent was 10-12-fold lower than the reaction with rRS.

**Figure 2.**
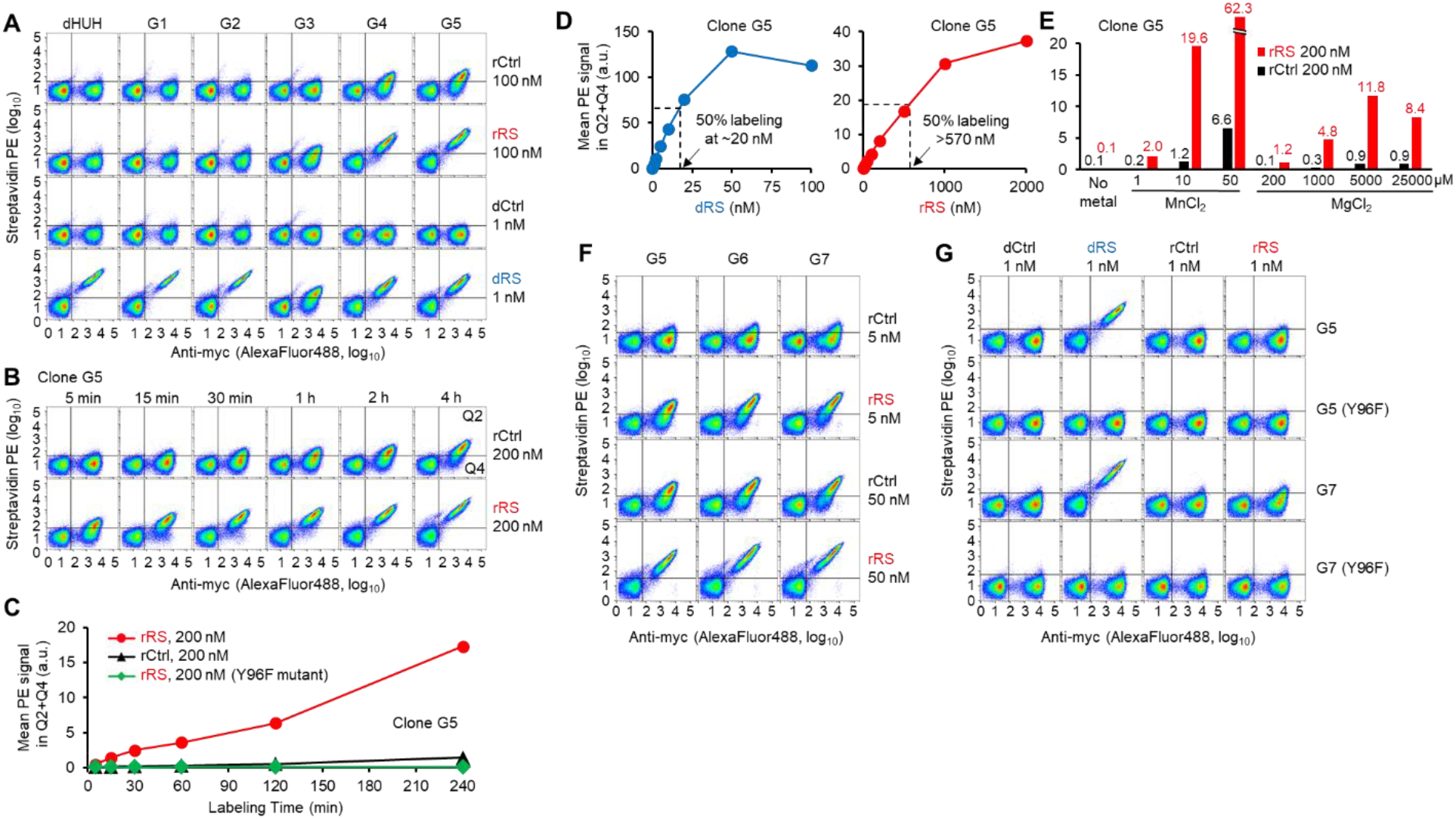
Directed evolution results and characterization of top clones on yeast. (**A**) FACS comparison of original dHUH template and evolved clones G1-G5, illustrating progress of evolution. Yeast displaying the indicated construct were labeled with rRS or dRS for 2 hours in the presence of 5 mM MgCl_2_ before streptavidin-PE/anti-myc staining and FACS analysis. rCtrl and dCtrl are scrambled versions of rRS and dRS, respectively (Figure 1D). (**B**) Time-dependent labeling of clone G5 on the yeast surface. 5 mM MgCl_2_ was present. (**C**) Quantification of data in (B) and for a negative control G5 mutant with the active site tyrosine ablated (Y96F, green). (**D**) Titration of DNA/RNA probe concentrations, with G5 on yeast. Labeling time was 5 min and 50 μM MnCl_2_ was present. FACS plots in Supplementary Figure 2A. (**E**) Effect of metal ion type and concentration on extent of G5 labeling on yeast with rRS probe (2 hours labeling). FACS plots in Supplementary Figure 2E. (**F**) Evaluation of clones G5-G7 on yeast at two different rRS probe concentrations, in the presence of 5 mM MgCl_2_. (**G**) Comparison of G5 and G7 on yeast with 1 nM rRS or dRS in the presence of 5 mM MgCl_2_. Negative controls with inactive Y96F mutants are also shown.

We performed more careful characterization of clone G5 (which has 9 mutations relative to dHUH). First, we performed a time course experiment, measuring the extent of labeling with 200 nM rRS from 5 minutes to 4 hrs. We observed a monotonic increase over this time, while the reaction with scrambled rRS rose much less (11-fold difference in labeling extent at 1 hour) (**Figures 2B-C**).

We next titrated the rRS (and for comparison, the dRS probe). **Figures 2D** and **S3A-C** show that G5 labeling with rRS increased steadily and did not show signs of saturation, even at the highest practically achievable concentration of 4 µM probe. By contrast, labeling with dRS could be easily saturated at ∼50 nM probe. Thus, G5 is still labeled with far greater sensitivity by DNA probe than by RNA probe. Because the HUH reaction mechanism requires a divalent metal ion - Mn^2+^ in the crystal structure - we tested the metal ion dependence of G5 labeling by rRS (**Figure 2E**). Like with wild-type dHUH, Mn^2+^ was preferred over Mg^2+^, but labeling was substantial at 1-5 mM of MgCl_2_, which is in the range of physiological Mg^2+^ concentrations (17, 18).

Mutagenesis of the active site Tyr96 in G5 to Phe abolished labeling by rRS, indicating that the labeling is of a covalent nature (**Figure S3D**). To examine the necessity of all 9 mutations in G5 relative to wild-type dHUH, we reverted each one to the wild-type amino acid identity. **Figure S4** shows that every reversion decreased reaction with dRS, with the mutations H22N and W82H having the largest effect. His82 in the wild-type structure forms a π-π stacking interaction with adenosine in the +1 position of dRS. His82 was mutated to Tyr in G3, and mutated further to Trp in G4, both of which may increase π-π stabilization. Mutation of Asn22 to the larger His sidechain in G5 would appear to introduce major clashes (with the U-1 nucleotide), unless rRS docks to G5 in a different conformation than dRS docks to dHUH.

Because of G5’s limited sensitivity and off-target activity, we performed two more generations of directed evolution, producing G6 and G7. For these selections, we decreased the concentration of rRS drastically, from 200 nM down to 1 nM. We also attempted to incorporate negative selections against scrambled probe (rCtrl) to improve the specificity of rHUH, but our efforts were not successful. As shown in Figure S5, we explored alternative paths in Generation 5 and in Generation 6 that utilized alternating positive and negative selections. Each time we implemented a negative selection, enriching low streptavidin-PE cells after treatment with 200-500 nM rCtrl, both the signal (rRS labeling activity) and background (rCtrl labeling activity) of the resulting library decreased. Both were then restored following rounds of positive selection. At the end of 5 alternating rounds, none of the enriched clones from this path had higher activity than the starting template (G5). We attempted the same strategy in Generation 7 (Supporting Figure S5C), and found that all of our enriched clones were identical in sequence to the template (G6). Ultimately the winners of each generation (G6 and G7) derived from the selection paths that largely did not incorporate negative rounds.

**Figure 2F** shows a comparison of G5, G6 and G7, demonstrating that G6 and G7 are indeed more reactive towards rRS than G5, although they also produce more signal with scrambled rCtrl. Across generations, the rRS/rCtrl signal ratio remains relatively constant, ∼11 fold for G5, G6, and G7. In **Figure 2G**, we compare G5 and G7 side by side at very low probe concentration – 1 nM rRS. Whereas labeling of G5 is difficult to detect at this concentration, G7 shows clear labeling over background. Negative controls with scrambled rCtrl or the inactivating point mutation Y96F in G7 fail to show labeling.

Altogether, our experiments indicate that our directed evolution has produced rHUH candidates (G5 and G7) with vastly increased reactivity towards RNA probes. Each generation of evolution produced steep benefits in reactivity, with G5 being ∼25-fold more active than G3, and G7 being ∼2.6-fold more active than G5. Ultimately, G7 could be labeled with as little as 1 nM rRS probe, in the presence of 5 mM Mg^2+^ rather than Mn^2+^, and with a specificity ratio of ∼12.

### In vitro characterization of rHUH

We purified our best rHUH clone, G7, by overexpression in *E. coli* and nickel affinity chromatography (16). Because rHUH has multiple surface-exposed histidines and a divalent metal ion binding site, it does not require a His6 tag to bind to nickel. However, we fused a myc tag to the C terminus to facilitate detection by Western blotting. To evaluate the reactivity of rHUH towards rRS, we used a gel shift assay, detecting the rHUH protein by its myc tag. Upon covalent reaction with rRS, the molecular weight of rHUH shifts up from ∼15 kD to ∼19 kD. Figure 3A shows the reaction of rHUH with both rRS and dRS, after just 1 minute incubation in the presence of 1 mM MnCl_2_. Mutation of the active site tyrosine (Y96F) abolishes labeling with both probes. Consistent with our observations on the yeast surface, rHUH also reacts significantly with scrambled control – both rCtrl and dCtrl, with a specificity ratio of 1.8-2.0 for RNA and 4.5-9.6 for DNA.

**Figure 3.**
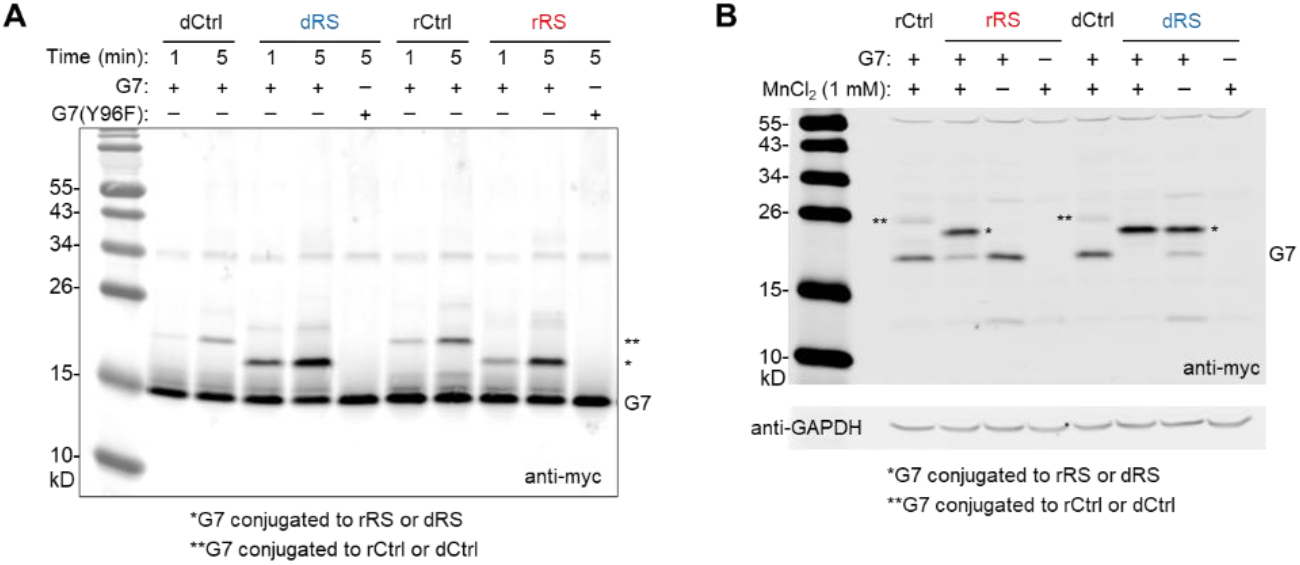
In vitro characterization of purified rHUH. (**A**) Gel shift assay showing reaction of purified rHUH (G7) with DNA and RNA probes. 0.5 µM G7 was combined with 10 µM rRS or dRS for 1 or 5 minutes in the presence of 1 mM MnCl_2_, then analyzed by SDS-PAGE and Western blotting with anti-myc antibody. (**B**) Same as (A) but rHUH (G7) was expressed in HEK 293T cells, and lysates were labeled with 10 µM rRS or dRS for 5 minutes in the presence of 1 mM MnCl_2_.

Next, we performed a more demanding experiment. We expressed rHUH (G7) in mammalian HEK 293T cells with a myc tag and using a TRE3G promoter. We then lysed the cells and treated the lysate with 10 µM of rRS or dRS for 5 minutes. Gel shift analysis in Figure 3B shows extensive reaction with both probes. Interestingly, rRS reaction requires supplementary addition of 1 mM MnCl_2_ while reaction with dRS does not. Here again, some non-specific reactions with rCtrl and dCtrl were detected, but these were attenuated compared to the previous experiment. The specificity ratios were 6.3 for RNA and 14.4 for DNA. Perhaps the presence of cellular components (proteins, other RNAs, etc) and lysis buffer (50 mM Tris-HCl, pH 7.5, 150 mM NaCl, 1% Nonidet P40) helps to reduce non-specific interactions between rHUH and non-cognate substrates.

Finally, we tested in vitro labeling of rHUH in the presence of high concentrations of cellular RNA. We prepared total RNA extract from HEK 293T cells and spiked it into 5-minute reactions of rHUH with rRS or rCtrl probe. Figure S6 shows that even a low concentration of exogenous RNA (0.1 µg/µL) suppresses labeling with rRS probe. This experiment highlights the challenge of transitioning this methodology to the interior of living cells.

## Discussion

In this study, we have engineered rHUH, a covalent and sequence-specific protein tag for RNA, through 7 generations of yeast surface display directed evolution. rHUH forms a covalent adduct with a 10 nucleotide RNA recognition sequence called rRS through Mn^2+^ or Mg^2+^-dependent formation of a tyrosine-phosphate ester bond. We showed that rHUH can couple to nonstructured RNAs on the yeast surface and in vitro, with a sensitivity down to 1 nM RNA probe, and a specificity factor of ∼6-12. This methodology could potentially be used to link both short and long RNA transcripts to a wide range of protein-based probes including fluorescent proteins, CRISPR-based editing enzymes, and proximity labeling tools.

rHUH enhances the arsenal of methods available to link RNA to protein. Among covalent strategies, chemical conjugation is dominant (19), but recent studies have begun to exploit enzymes for the task. For example, the RNA-TAG method (20) uses a tRNA guanine transglycosylase to attach a benzylguanine moiety onto an RNA of interest. After purification of the conjugate, the RNA is reacted with a SNAP tag-fused protein for 4 hours (20). A simpler methodology uses an engineered variant of uridine-54 tRNA methyltransferase, TrmA, to form a covalent adduct with an RNA hairpin motif (21), but the reaction appears to be slow and the generalizability has not been explored.

The long-term goal of rHUH engineering is protein-RNA conjugation inside living cells, where the ability to visualize, analyze, and manipulate specific RNA species would be very highly enabling. The major challenges to translating rHUH methodology to the cell interior are sensitivity, specificity, and metal ion availability. While rRNAs and tRNAs are highly abundant (∼2-25 uM (22)), many mRNA species are present at just 1-100 nM, or even less, inside cells (22). An effective rHUH tag would need to react rapidly with target RNA species in this concentration regime, while avoiding off-target reactions with thousands of higher-abundance RNAs. The current specificity ratio of our rHUH (ratio of reaction with rRS versus scrambled control is only 6-12, far less than what is required for specific tagging in living cells. Finally, wild-type dHUH requires 10-500 µM of Mn^2+^, which is scarce inside cells; protein-bound and unbound pools of Mn^2+^ total only ∼1 µM (23)). We used directed evolution to shift the divalent metal ion requirement towards Mg^2+^, which is present at 0.5-1 mM in free form (18), but further engineering is likely necessary for high activity in the cellular milieu.

Our yeast display directed evolution platform was able to produce RNA labeling activity from a starting template with no detectable reactivity towards RNA on the yeast surface. We observed remarkable improvements in activity towards RNA over 7 generations of directed evolution, culminating in our best clone, G7, which is ∼780 fold more active than G1, our first-generation winner. Nevertheless, we found that our selection platform was highly limited when it came to engineering rHUH sequence-specificity. We attempted many rounds of negative selection but only managed to decrease *both* specific and non-specific reactivity together. From generation 3 onward, only the activity improved, with little to no gain in specificity ratio. A different selection scheme, that simultaneously (rather than alternately) selects for high rRS labeling and minimal off-target labeling might yield improved results. Our study lays the groundwork for future improvements in this new tool class for investigating the biology of RNAs.

## Methods

### Cloning

The double-digested vectors and the PCR amplified inserts with 20-50 homologous overhangs on both ends were ligated by Gibson assembly. Ligated plasmid products were introduced by heat shock transformation into competent XL1-Blue bacteria.

### Yeast cell culture

For yeast-display, S. cerevisiae strain EBY100 cultured in yeast extract peptone dextrose (YPD) complete medium was transformed with the yeast-display plasmid pCTCON2 (24) using the Frozen E-Z Yeast Transformation II kit (Zymo Research) according to manufacturer protocols. Transformed cells were selected on synthetic dextrose plus casein amino acid (SDCAA) plates and propagated in SDCAA medium at 30 °C. Protein expression was induced by inoculating saturated yeast culture into SD/RCAA (SDCAA medium with 90% of dextrose replaced with raffinose) and incubated at room temperature for 4 h, then 2% of galactose was added to the culture and continue incubating at room temperature with shaking overnight.

### Generation of rHUH libraries for yeast display

Libraries of PCV2 mutants were generated by error-prone PCR according to published protocols (25). 100 ng of the template in vector pCTCON2 was amplified for 20 cycles with 0.4 µM forward and reverse primers:

F: 5’-CTAGTGGTGGAGGAGGCTCTGGTGGAGGCGGTAGCGGAGGCGGAGGG TCGGCTAGC-3’

R: 5’-TATCAGATCTCGAGCTATTACAAGTCCTCTTCAGAAATAAGCTTTTGTTC

GGATCC-3’ and 1x ThermoPol Reaction Buffer (NEB), 10 units of Taq DNA polymerase (NEB), 5, 10, or 20 µM of 8-oxo-2’-deoxyguanosin-5’-triphosphate (8-oxo-dGTP), 1, 2, or 4 µM of 2’deoxy-P-nucleoside-5’-triphosphate (dPTP), 200 µM of dNTP in a 100 µl total volume. 300 ng of the gel purified PCR products were reamplified for another 30 cycles under normal PCR conditions using following the primer pair:

F: 5’-CAAGGTCTGCAGGCTAGTGGTGGAGGAGGCTCTGGTG-3’

R: 5’-CTACACTGTTGTTATCAGATCTCGAGCTATTACAAGTC-3’

The amplified PCR products were gel purified and combined with BamHI-NheI linearized pCTCON2 vector (4 µg insert/1 µg vector) and electroporated into electrocompetent S. cerevisiae EBY100 (25). The electroporated cultures were rescued in 2 ml YPD medium for 1 h at 30 °C with no shaking. The rescued cell suspension was transferred to 100 mL of SDCAA medium supplemented with 50 units/ml penicillin and 50 μg/ml streptomycin and grown for 2 days at 30 °C.

### Biotinylated dRS and rRS probes

All biotinylated probes were ordered from IDT with 3’ Biotin-TEG modifications. All probes were received as dried pellets and dissolved to 1 mM stocks with H_2_O upon arrival. A fraction of 1 mM stocks was further diluted with H_2_O into 100 µM secondary stocks. The secondary stocks were aliquoted and frozen under -80 °C. The 1 mM stocks were frozen under -80 °C in the original tube.

### General method for labeling of yeast-displayed rHUH variants

All reagents used in rHUH labeling are in RNase-free format or as clean as possible. Yeast cells induced overnight were washed twice with PBST buffer (PBS, pH 7.4, supplemented with 0.2% Tween-20) and resuspended in PBST supplemented with 10 µM MnCl2 or 5 mM MgCl2 (or indicated otherwise) and RNase inhibitor cocktail (RiboLock, Invitrogen). The indicated amount of probe was added to the cells and incubated at room temperature with gentle shaking. After labeling, the yeast cells were washed once with PBST supplemented with 5 mM EDTA, followed by incubation with anti-myc chicken IgY antibody (Exalpha Biological, 1:400) and then with streptavidin-PE (Jackson ImmunoResearch, 1:400) and Goat anti-chicken IgY antibody AlexFlour 488 (AF488) conjugate (Invitrogen, 1:400). The labeled yeast cells were analyzed by ZE5 FACS cell analyzer (Bio-Rad).

### General methods for yeast display-based directed evolution

For the first round of selection for each generation, the electroporated yeast libraries were grown for 2 days to reach saturation. Then ∼1.5×109 of yeast cells from the saturated culture were inoculated into 165 ml of SD/RCAA medium and incubated at room temperature for 4 h, then 2% of galactose was added to the culture and continue incubating at room temperature with shaking overnight. ∼6×108 of yeast cells from the overnight induced culture were labeled following the method described in section “General methods for yeast displayed rHUH labeling”. The labeled yeast cells were resuspended in 10 ml of ice-cold PBS and sorted on a BD FACS Aria II cell sorter (BD Biosciences) with appropriate lasers and emission filters for PE and AF488. The yeast cells were processed at a rate of ∼20,000 cells per second and the top 0.5-1% of cells with highest PE and AF488 signals were collected in SDCAA medium containing 1% penicillin-streptomycin and incubated at 30 °C for 3 days. Limited by the sorting speed and FACS sorter availability, ∼3×108 of yeast cells were processed by the sorter and ∼2×106 of yeast cells were collected.

For the second round, ∼2×107 of yeast cells were labeled and sorted following similar procedures. For the third and later rounds, ∼4×106 of yeast cells were used. For positive selections, the top 0.5-1% of cells with highest PE and AF488 signals were collected. For negative selections, the top 1% of cells with highest AF488 signal and lowest PE signal were collected.

After the last round of selection, the collected yeast cells were grown to saturation and 1 ml of saturated culture was removed for DNA extraction using the Zymoprep yeast Plasmid Miniprep II (Zymo Research) kit according to manufacturer protocols (using 10 µl zymolyase). The extracted DNA was transformed into competent XL1-Blue bacteria. After the emergence of colonies on the bacteria culture plate, 12-24 colonies were picked up subjected to standard plasmid amplification protocols. The amplified plasmids were analyzed by Sanger sequencing.

### Mammalian cell culture and transfection

HEK293T cells from ATCC (passage number <25) were cultured as a monolayer in growth media (DMEM, high glucose, Gibco) supplemented with 10% fetal bovine serum (Avantor) at 37 °C under 5% CO2. For transient expression, cells were typically transfected at approximately 80% confluency using 1 mg/ml PEI max solution (polyethylenimine HCl max pH 7.3). For inducing expression or rHUH, 100 µg/ml of doxycycline was added to the medium.

### Bacterial expression and purification of rHUH protein

For bacterial expression and purification of rHUH protein, we cloned the rHUH sequence into the pTD68 vector (Addgene 123643) with all N-terminal tags removed and a C-terminal myc tag added. We found that rHUH from G5 onward could bind to Ni-NTA without a His6 tag. Competent BL21 E. coli was transformed with the rHUH expression plasmid by heat shock. Cells were then grown in LB medium containing 100 mg/L carbenicillin at 37 °C and 220 r.p.m. until an OD600 of ∼0.6-0.8 was reached. Protein expression was induced with 0.05 mM isopropyl β-D-1-thiogalactopyranoside (IPTG), then the culture was shifted from 37 °C to room temperature during the induction period. After overnight growth, the bacteria were pelleted by centrifugation at 7,000 ×g for 3 min at room temperature, the supernatant was discarded, and the pellet was stored at -80 °C.

The frozen pellet was thawed at room temperature and transferred to ice immediately after thawed. 10 ml of lysis buffer (B-PER Bacterial Protein Extraction Reagent, Thermo Scientific, supplemented with 1% protease inhibitor cocktail and 1 mM PMSF) per 500 ml of culture was added to the pellet. The pellet was resuspended and kept on ice for at least 10 min. Then the lysate was sonicated using a Misonix sonicator (1-s on, 1-s off, for a total of 60-s on, ice chill for 2 min. Repeat 4 times). The sonicated lysate was clarified by centrifugation for 10 min at 18,000 ×g at 4 °C, and the supernatant was clarified by same method again, and then the supernatant was carefully transferred to a 10-ml conical with 2 ml Ni-NTA agarose bead slurry (Invitrogen) per 10 ml lysate and incubated at 4 °C for 30 min with gentle rotation. Then the slurry was placed in a gravity column and washed twice with 10 CV of washing buffer (50 mM Tris-HCl, pH 8.0, 500 mM NaCl, 30 mM imidazole). The protein was eluted with 5 CV of elution buffer (50 mM Tris-HCl, pH 8.0, 500 mM NaCl, 300 mM imidazole). The purity was analyzed by Coomassie Blue staining.

The eluted protein was further purified by using a HiLoad 16/600 Superdex 75 pg column (Cytiva) with FPLC buffer (50 mM Tris-HCl, pH 7.5, 300 mM NaCl, 1 mM EDTA, 1 mM DTT). The collected fractions were concentrated using Amicon Ultra-15 Centrifugal Filter Units, 3,000-kDa cutoff.

### In vitro labeling of purified rHUH or rHUH-containing cell lysate and Western blot analysis

Purified rHUH proteins were diluted in 50 mM Tris-HCl, pH 7.5, 50 mM NaCl, and 1 mM MnCl2 (or otherwise indicated). DNA/RNA probes were added and mixed by a brief vortex. One fifth volume of 6× SDS Protein Loading buffer supplemented with 5 mM EDTA was added to quench the reaction. And the reaction was further quenched by heat denaturation at 95 °C for 5 minutes. For HEK293T cells expressing rHUH, the cells were lysed with Nonidet P40 (NP40) lysis buffer (50 mM Tris-HCl, pH 7.5, 150 mM NaCl, 1% NP40) on ice for 10 min. The lysate was clarified by centrifugation for 10 min at 20,000 ×g at 4 °C. The supernatant was transferred to a new tube and mixed with 1 mM MnCl2 and DNA/RNA probes. After the reaction, one fifth volume of 6× SDS Protein Loading buffer supplemented with 5 mM EDTA was added to quench the reaction. And the reaction was further quenched by heat denaturation at 95 °C for 5 minutes.

The denatured mixtures were resolved by a 15% Tris-Glycine gel with 10% SDS, and then transferred to a nitrocellulose membrane. The membrane was briefly washed by TBST (100 mM Tris-HCl, pH 7.5, NaCl 150 mM, 0.05% Tween-20) followed by incubation with 5% non-fat milk (Lab Scientific bioKEMIX) in TBST for 1 h. Then the membrane was incubated with anti-myc chicken IgY antibody (Exalpha Biological, 1:5,000, in TBST) at 4 °C overnight. After three washes with TBST, the membrane was incubated with IRDye 680RD Donkey anti-Chicken secondary antibody (LI-COR Biosciences, 1:5,000, in TBST, protected from light) for 1 h. Then the membrane was washed three times with TBST and imaged by a LI-COR Odyssey CLx gel imager.

## Supporting Figures

**Supporting Figure 1.**
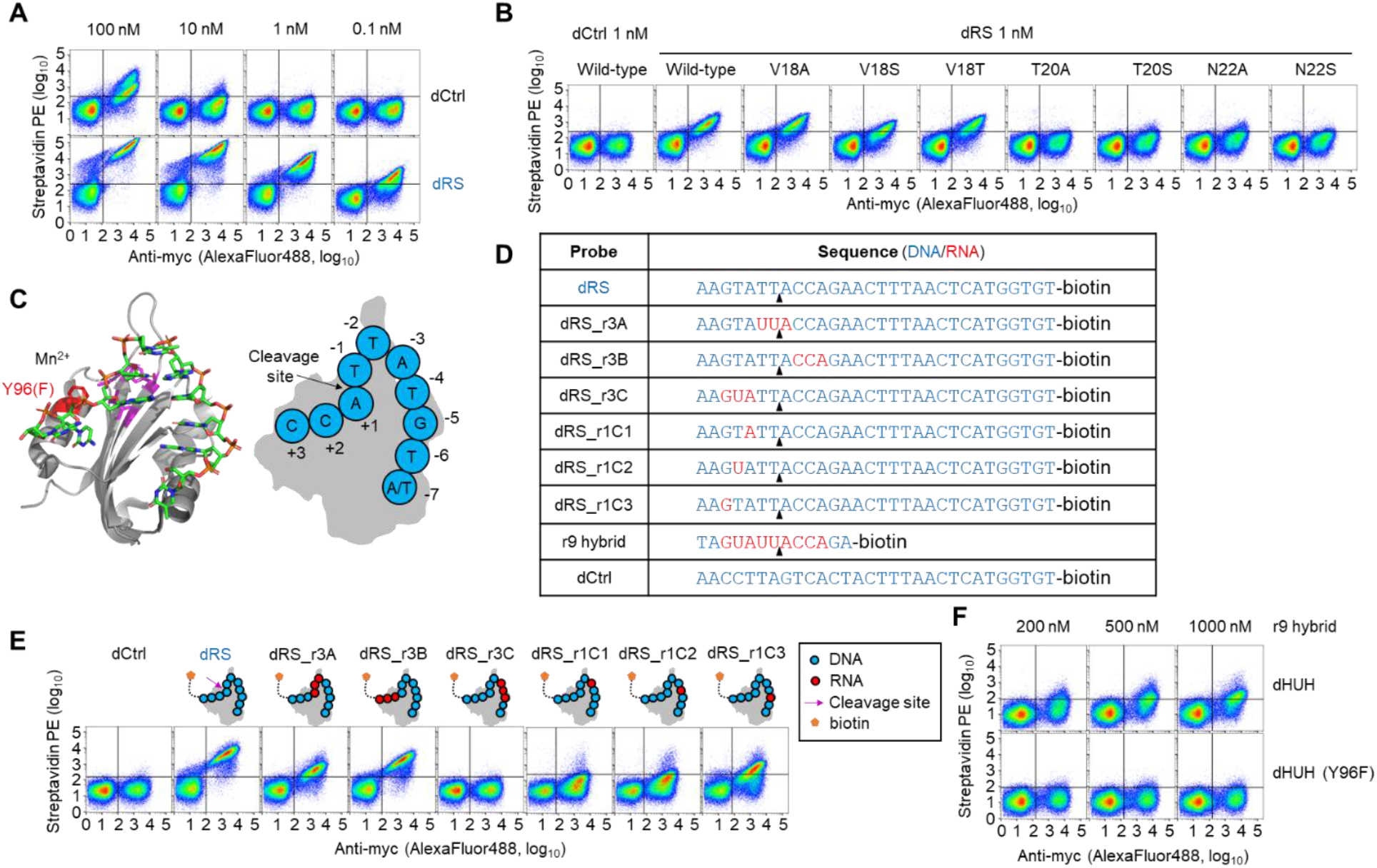
Initial evaluation of dHUH point mutants and RNA-DNA hybrid probes on the yeast surface. (**A**) Labeling of wild-type dHUH on yeast with various concentrations of dRS probe. (**B**) Point mutants of dHUH affect labeling with 1 nM dRS (10 minutes reaction time with 10 µM MnCl_2_). (**C**) Structure of wild-type dHUH in complex with dRS (PBD ID 6WDZ (16)). In cartoon at right, nucleotides of dRS are shown as blue circles, and labeled with their positions relative to the cleavage site. (**D**) Table of RNA-DNA hybrid probes tested. (**E**) FACS analysis of wild-type dHUH-displaying yeast cells, after reaction with hybrid probes (at 1 nM for 15 min in the presence of 10 µM MnCl_2_). Positions -3 and -4 of dRS are most sensitive to alteration. (**F**) FACS analysis of wild-type dHUH-displaying yeast cells, after reaction with r9 hybrid probe at different concentrations for 2 hours in the presence of 10 µM MnCl_2_.

**Supporting Figure 2.**
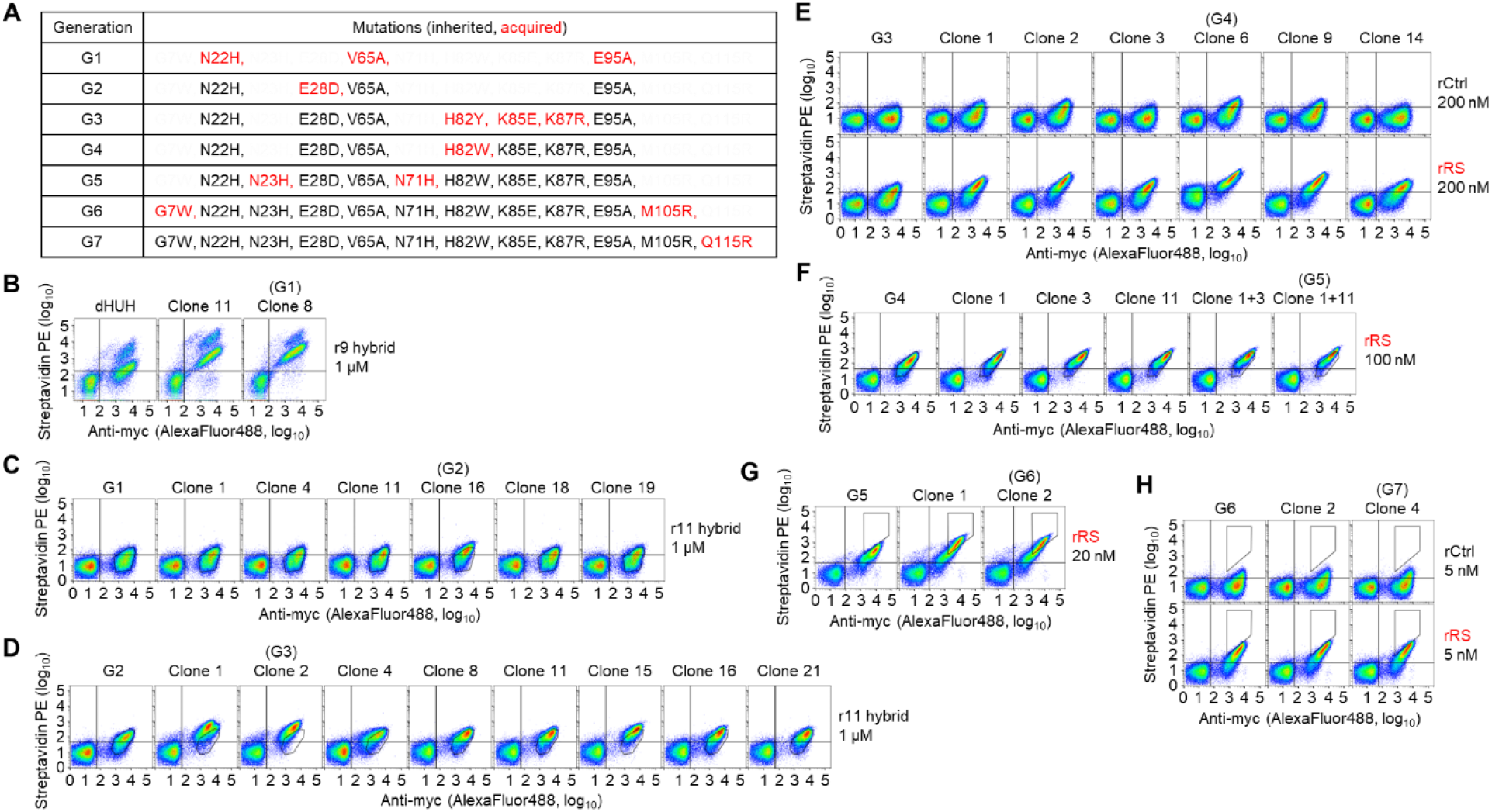
Sequences and characterization of enriched clones on yeast. (**A**) Summary of mutations found in clones G1-G7. (**B-H**) Analysis of enriched clones from Generations 1-7. All reaction times were 2 hours. In (B-E), 10 uM of MnCl_2_ was used. In (F-H), 5 mM of MgCl_2_ was used. In (**F**), Clone x+y means that mutations in x and y are combined together.

**Supporting Figure 3.**
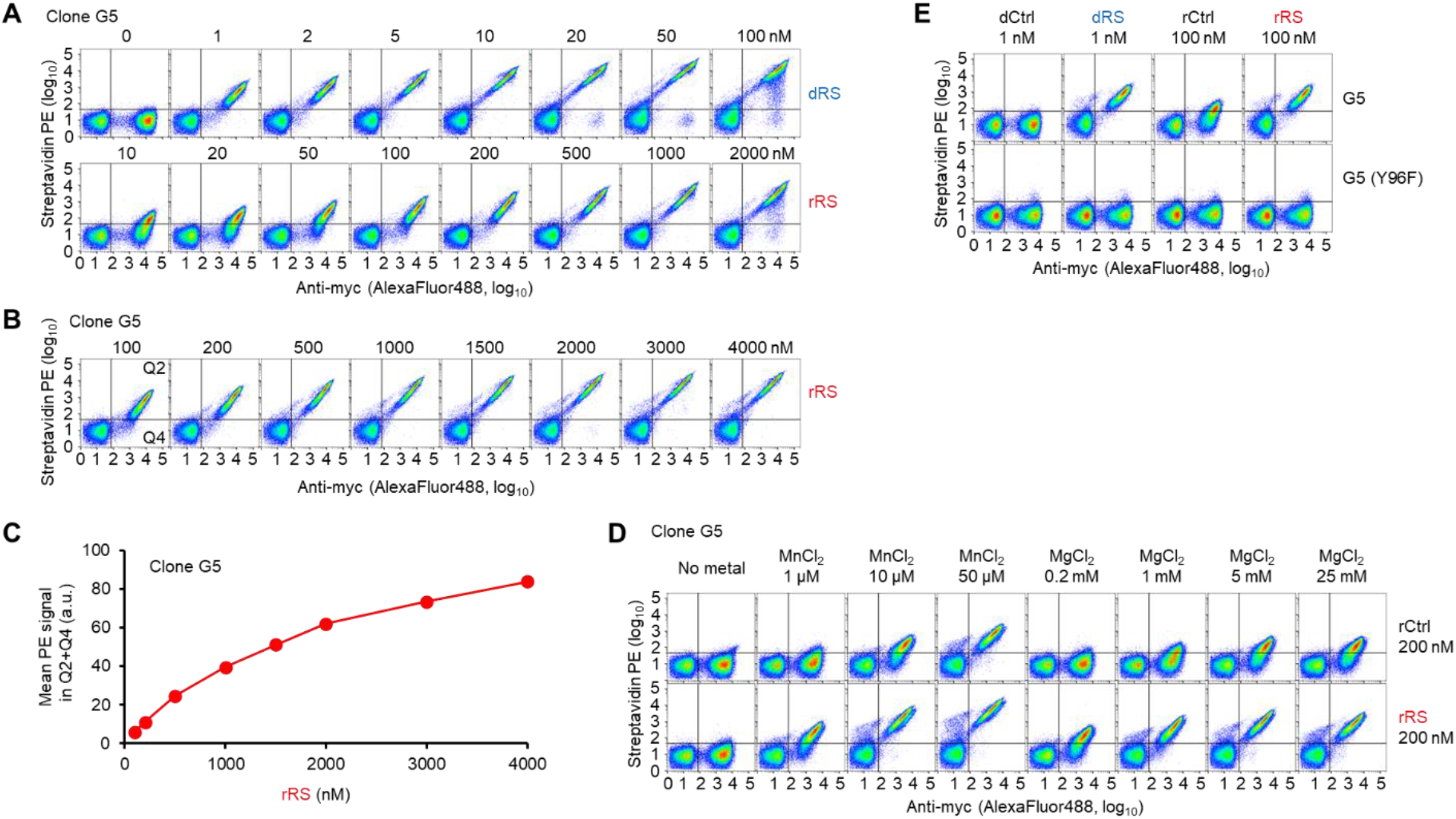
Additional characterization of clone G5 on yeast. (**A**) FACS plots for substrate concentration titration shown in **Figure 2D**. (**B**) Titration to even higher rRS probe concentrations with G5 yeast. (**C**) Quantification of data in (B). (**D**) FACS plots for metal ion analysis in **Figure 2E**. (**E**) Testing inactive point mutant (Y96F) of G5 on yeast at high probe concentration (100 nM).

**Supporting Figure 4.**
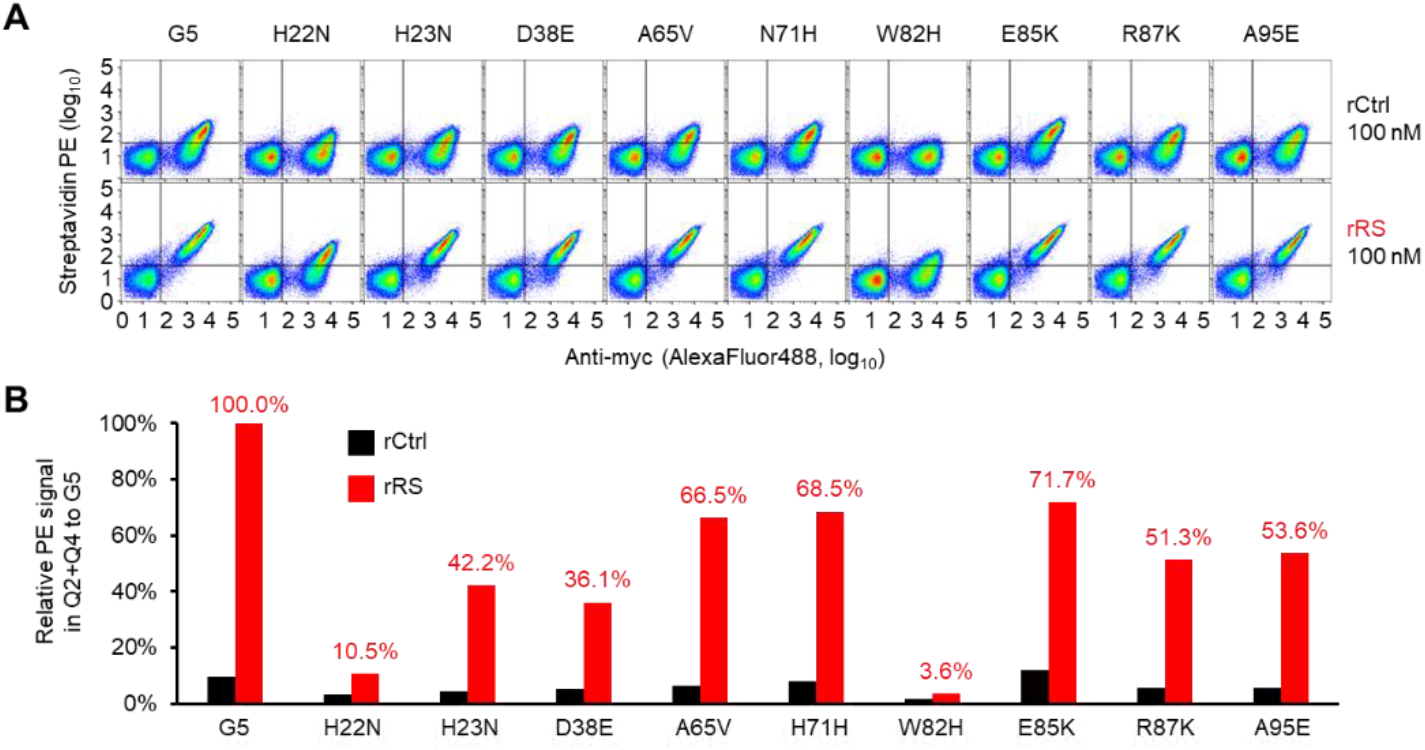
Evaluating the necessity of mutations in Clone G5. (**A**) All of the 9 mutations in G5 were individually reverted to the wild-type amino acid, and tested on the yeast surface with 100 nM rRS for 2 hours in the presence of 5 mM MgCl_2_. (**B**) Quantification of data in (A). Percentage of active (streptavidin-PE+) cells normalized to that of G5.

**Supporting Figure 5.**
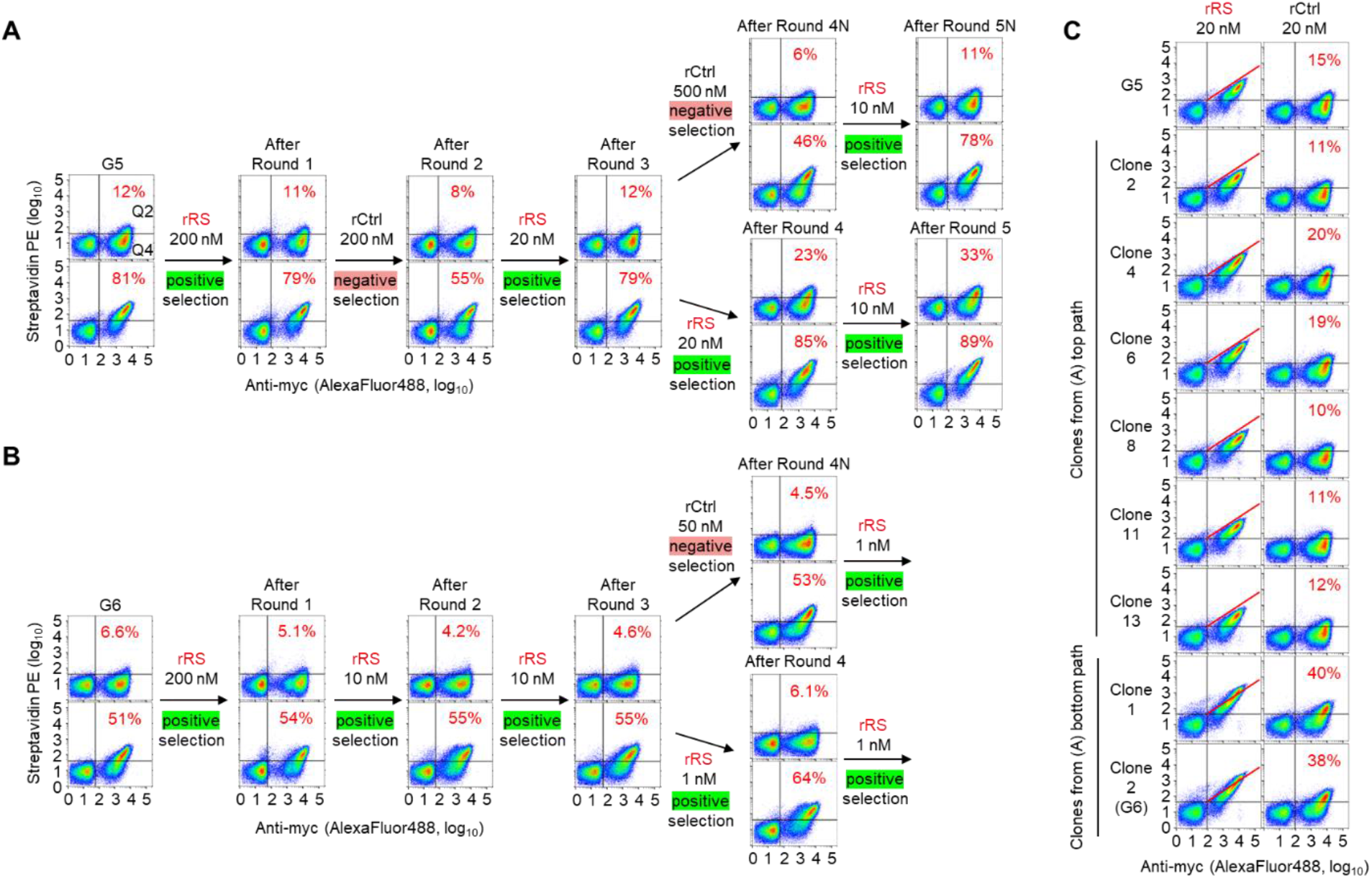
Alternate selection paths used in Generations 6 and 7 incorporation negative selection rounds. (**A**) Alternate generation 6 path, starting from G5 as template. Top plots show reaction with 20 nM rCtrl; bottom plots show reaction with 20 nM rRS. (**B**) Alternate generation 7 path, starting from G6 as template. Top plots show reaction with 5 nM rCtrl; bottom plots show reaction with 5 nM rRS. (**C**) Analysis of clones enriched from (A). For these analyses, all samples were labeled for 2 hours at room temperature in the presence of 5 mM MnCl_2_. Percentages calculated as fraction of cells in Q2 divided by cells in Q2+Q4.

**Supporting Figure 6.**
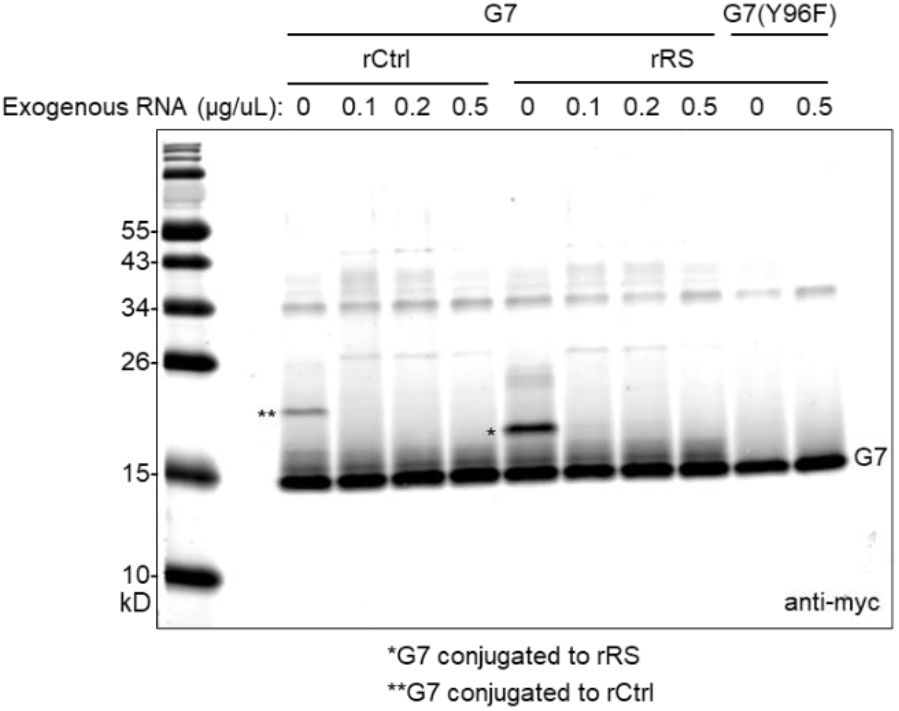
In vitro labeling of rHUH is suppressed by addition of exogenous RNA. Gel shift assay showing reaction of purified rHUH (G7) with rRS, in the presence versus absence of total RNA extract from HEK 293T cells. 0.5 µM G7 was combined with 10 µM rRS or rCtrl for 5 minutes in the presence of 0-0.5 µg/µL RNA extract from HEK 293T cells. Samples were analyzed by SDS-PAGE and Western blotting with anti-myc antibody.

